# A toxic environment selects for specialist microbiome in poison frogs

**DOI:** 10.1101/2024.01.10.574901

**Authors:** Stephanie N. Caty, Aurora Alvarez-Buylla, Cooper Vasek, Elicio E. Tapia, Nora A. Martin, Theresa McLaughlin, Peter K. Weber, Xavier Mayali, Luis A. Coloma, Megan M. Morris, Lauren A. O’Connell

## Abstract

Shifts in microbiome community composition can have large effects on host health. It is therefore important to understand how perturbations, like those caused by the introduction of exogenous chemicals, modulate microbiome community composition. In poison frogs within the family Dendrobatidae, the skin microbiome is exposed to the alkaloids that the frogs sequester from their diet and use for defense. Given the demonstrated antimicrobial effects of these poison frog alkaloids, these compounds may be structuring the skin microbial community. To test this, we first characterized microbial communities from chemically defended and closely related non-defended frogs from Ecuador. Then we conducted a laboratory experiment to monitor the effect of the alkaloid decahydroquinoline (DHQ) on the microbiome of a single frog species. In both the field and lab experiments, we found that alkaloid-exposed microbiomes are more species rich and phylogenetically diverse, with an increase in rare taxa. To better understand the strain-specific behavior in response to alkaloids, we cultured microbial strains from poison frog skin and found the majority of strains exhibited either enhanced growth or were not impacted by the addition of DHQ. Additionally, stable isotope tracing coupled to nanoSIMS suggests that some of these strains are able to metabolize DHQ. Taken together, these data suggest that poison frog chemical defenses open new niches for skin-associated microbes with specific adaptations, including the likely metabolism of alkaloids, that enable their survival in this toxic environment. This work helps expand our understanding of how exposure to exogenous compounds like alkaloids can impact host microbiomes.

## Introduction

The microbiota plays a role in many host biological processes, including development, behavior, and immunity ^1–3^. When a community of microbes interacts with a host, shifts in the microbial community structure can have large impacts on host health ^4–15^. One way that host-microbe interactions can be altered is through the addition of exogenous compounds to the system. For example, antibiotic treatment can lead to a depletion in abundance of one or more bacterial taxa, altering bacterial succession dynamics ^16^. However, hosts are also exposed to other exogenous compounds, the impacts of which are less clear. For example, while microbial physiology of the human gut is dramatically impacted by antibiotic treatment, this is not the case for some other drugs, including the cardiac glycosides digoxin and digitoxin ^17^. Given the significance of the microbiota for host health across many taxa, it is important to understand how various types of exogenous compounds alter host-associated microbial communities. Alkaloids are compounds of interest given their widespread ubiquity across ecosystems and the relative paucity of studies about their impacts on microbial communities.

Poison frogs (Family Dendrobatidae) offer a unique approach to study the interaction of microbiomes and exogenous chemicals because many species within this family consume and sequester xenobiotic alkaloids as part of their normal diet of small arthropods. Within the dendrobatid family, there are also species that do not sequester alkaloids and therefore are non-toxic, enabling comparisons of microbial communities from closely related frogs that differ in their alkaloid loads. For poison frogs that do sequester alkaloids, the compounds are sequestered to, stored, and secreted from granular glands in the skin for chemical defenses ^18–20^. This bioaccumulation of dietary alkaloids puts the frog skin microbial communities in contact with high concentrations of the alkaloids. Since the frogs do not synthesize their own alkaloids, but instead obtain them from their diet, frogs are non-toxic in the laboratory. Laboratory reared frogs can also be fed alkaloids experimentally to recapitulate the toxic phenotype ^21–23^. Our ability to rear poison frogs in the lab enables comparisons across lab and field studies to infer generalizable principles of chemical defense and microbial community dynamics.

Many naturally occurring alkaloids are antimicrobial ^24–27^, including those found on the skins of poison frogs. Alkaloid cocktails taken from the Strawberry poison frog (*Oophaga pumilio*) deter the growth of common lab strains of bacteria (*Escherichia coli*) and fungi (*Candida albicans*), and common amphibian bacterial pathogens, including *Klebsiella pneumoniae* and *Aeromonas hydrophila* ^28–30^. In addition to a combination of alkaloids taken directly from frog skins, individual poison frog alkaloids vary in the strength and efficacy in deterring microbial growth ^28^. For example, some decahydroquinoline (DHQ) are active against *Bacillus subtilis*, while others are active against *C. albicans*, and no DHQs tested deterred *E. coli*. Importantly, these studies have been done with bacteria and fungi that do not belong to the typical frog microbiome, and to our knowledge, no studies have looked at the impact of alkaloids on individual members of the commensal skin microbiome. Moreover, only one study has described the composition of a poison frog microbiome ^31^, but this was in a single species of chemically defended frog, limiting the ability to identify alkaloid-microbe interactions. More in-depth characterizations of poison frog microbiome-alkaloid interactions can help to elucidate more generalizable patterns in how alkaloids shape commensal microbial communities.

The goal of this study was to examine how alkaloids shape the microbial communities found on the skins of poison frogs and to identify mechanisms that may explain these community-level changes. We hypothesized that alkaloid exposure would result in less diverse microbial communities given the previously demonstrated antimicrobial properties of many poison frog alkaloids. Furthermore, we anticipated that the microbes that could survive in this habitat would exhibit specific adaptations that enable persistence in this toxic environment.

## Results

### Frogs with high alkaloid loads had more diverse but less even microbial communities

To determine how alkaloids correlate with the microbial communities of poison frogs, the bacterial and fungal communities of frog skins were characterized for 11 species of frogs across 9 geographic locations in Ecuador (Fig 1A). The frogs can be grouped into three categories of summed alkaloid load as defined by the summed area under the curve of all alkaloids: high, low, and no alkaloids (p <0.001 for all pairwise comparisons; Fig 1B). In both bacterial and fungal communities, samples primarily clustered based on collection location as opposed to the species of frog (Figure 1C). The fungal and bacterial dendrograms were topologically similar to one another, with an entanglement score of 0.1, where 0 is completely identical and 1 is no overlap (Figure 1C). In a principal coordinates analysis (PCoA) using Bray-Curtis dissimilarity, the individual frog samples clustered primarily based on the location of the frog (PERMANOVA bacteria R^2^= 0.24, p <0.001; fungal R^2^ = 0.23, p <0.001), and secondarily by the species of frog (PERMANOVA bacteria R^2^= 0.19, p <0.001; fungal R^2^=0.14, p <0.001). However, the alkaloid level of the frog also explained some of the variance in community composition, although less so than host species or location (PERMANOVA bacteria: Fig 1D, R^2^ = 0.03, p < 0.001, fungi: Fig 2A, R^2^ = 0.03, p <0.001). Overall, frog skin microbial community composition was influenced primarily by geographic location from which the host was collected and host species identity, but also by host alkaloid level.

**Figure 1.**
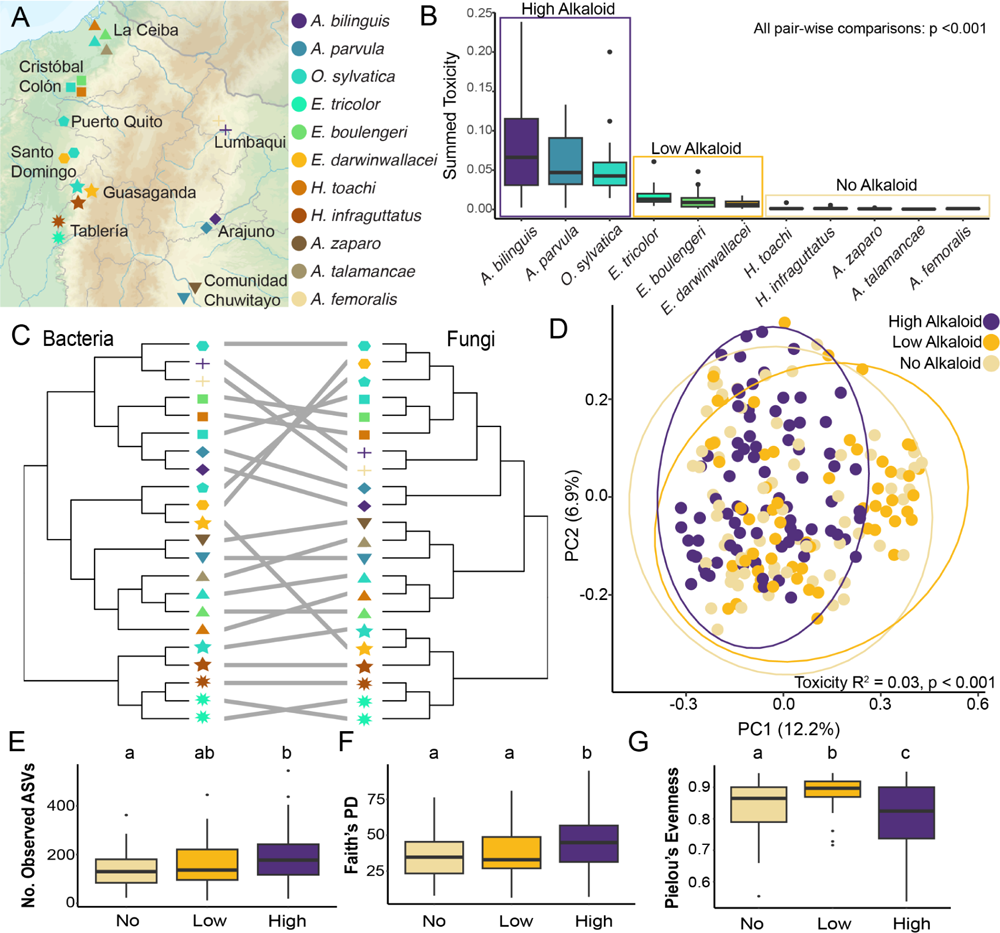
Bacterial communities are more diverse but less even in chemically defended frogs. **(A)** Eleven species of frogs (distinguished by color) were collected from nine locations (distinguished by shape) in Ecuador. **(B)** Based on summed alkaloid content, frog species can be grouped into high alkaloid quantity, low alkaloid quantity, and no alkaloids. Using a Kruskall-Wallace and post-hoc Dunn’s test, all pairwise comparisons have p <0.001. **(C)** Bacterial and fungal communities cluster primarily based on host collection location (shape) and exhibit similar clustering patterns (entanglement = 0.1). **(D)** High, low, and no alkaloid bacterial communities differ based on Bray-Curtis Dissimilarity. Bacterial communities from high alkaloid frogs are more speciose **(E)**, more phylogenetically diverse (**F**; PD = phylogenetic diversity), and less even **(G)**. Letters denote statistical differences based on Dunn’s post-hoc test with p value <0.05, dots are outlier points.

**Figure 2.**
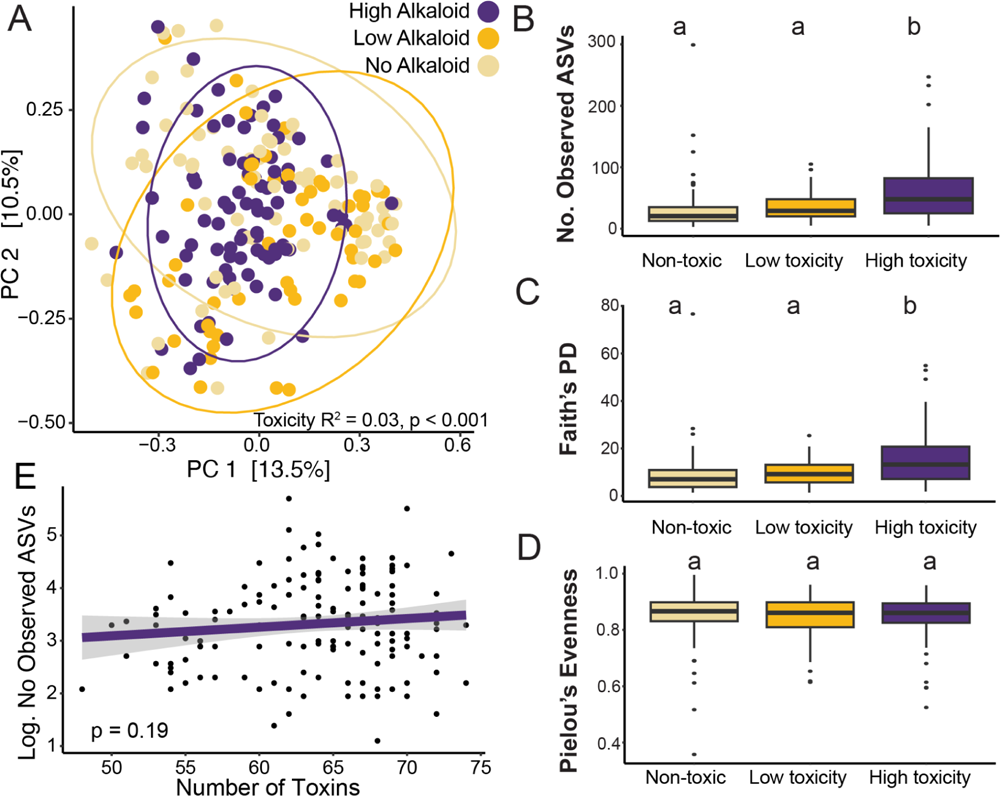
Alkaloid quantity shapes fungal communities and makes communities more diverse. **(A)** High, low, and no alkaloid fungal communities differ based on Bray-Curtis Dissimilarity. Fungal communities from high alkaloid frogs are more speciose **(B)** and more phylogenetically diverse (**C**; PD = phylogenetic diversity), but do not differ in evenness **(D)**. Letters denote statistical differences based on Dunn’s post-hoc test with p value <0.05, dots are outlier points. **(E)** Number of individual toxins found on a frog does not correlate with log transformed fungal ASV counts.

Chemically defended frogs had more diverse bacterial and fungal communities than undefended frogs. Specifically, high alkaloid frogs had a greater number of bacterial amplicon sequence variants (ASVs) than non-toxic frogs (Fig 1E, high vs. no alkaloids p = 0.002, other comparisons non-significant) and a greater number of fungal ASVs than either low alkaloid or no-alkaloid frogs (fungi: Fig 2B, high vs. low p = 0.04, high vs. non p <0.001). Furthermore, the phylogenetic diversity (as measured by Faith’s Phylogenetic Diversity) was higher in high alkaloid frogs than in either low alkaloid or no alkaloid frogs (bacteria: Fig 1F, high vs. low p = 0.02, high vs. non p = 0.001; fungi: Fig 2C, high vs. low p = 0.02, high vs non p <0.001). However, high alkaloid bacterial communities were less even than either low alkaloid or no-alkaloid frogs, suggesting the communities were dominated by particular taxa (Pielou’s evenness, bacteria: Figure 1G, p<0.001 for all pairwise comparisons). This was not the case for fungi, where communities across all alkaloid levels had the same evenness (Fig 2D, p = 0.7). There was no correlation between the toxin dataset and the bacterial ASV dataset (bacteria Mantel test p=0.63, fungi Mantel test p = 0.244). Furthermore, the number of individual alkaloids did not correspond with increased bacterial or fungal diversity (fungi: p = 0.19, Fig 2E; bacteria: p = 0.82, Supplementary Fig 1). These results show that the quantity of alkaloids, rather than type or diversity, correlated with the richness and evenness of microbial communities on poison frog skin.

The majority of differentially abundant taxa between alkaloid load groups were low abundance taxa. We identified taxa (agglomerated at the genus level) with structural zeros, which are taxa that are absent from a sample after statistical adjustment for sample size, in essence creating a presence/absence comparison of genera across groups. When comparing high alkaloid frogs to no alkaloid frogs, there were 124 bacteria genera and 78 fungal genera that were only present in high alkaloid frogs, while only 38 bacterial genera and 4 fungal genera were present only in no alkaloid frogs (Supplementary Table 1). These high-alkaloid only taxa comprised 5% of the total bacterial and 13% of total fungal communities of high alkaloid frogs, suggesting that some of these differences in diversity and community structure were coming from the addition of new, rare taxa in high-alkaloid frogs. Overall, we observed that alkaloid load played a significant role in shaping the microbial communities of wild frogs, and that, contrary to our expectations, high alkaloid loads increase microbial community diversity.

### A single alkaloid changed the resident skin microbial community

To directly test the hypothesis that alkaloids shift microbial communities and remove the confounding factors of environment and host species, we performed a controlled toxin feeding experiment with the Diablito poison frog (*Oophaga sylvatica*), a species included in our field data set capable of acquiring alkaloids from its diet. The alkaloid DHQ was selected because it is commercially available, and this class of alkaloids is found widely distributed across poison frog species (Supplementary Figure 2). Frogs were swabbed for five days prior to alkaloid administration, and for the 10 days of administration of the experimental treatment (DHQ) or vehicle control (water) (Fig 3A). Swabs were used for characterization of the bacterial and fungal communities with 16S and ITS amplicon sequencing. At the end of the feeding experiment, frog skins were used to quantify alkaloid load to confirm the deposition of DHQ on the skin. As expected, the *O. sylvatica* fed DHQ sequestered the DHQ onto their skin, while those without DHQ feeding had no DHQ on their skin (Fig 3B, p = 0.004).

**Figure 3.**
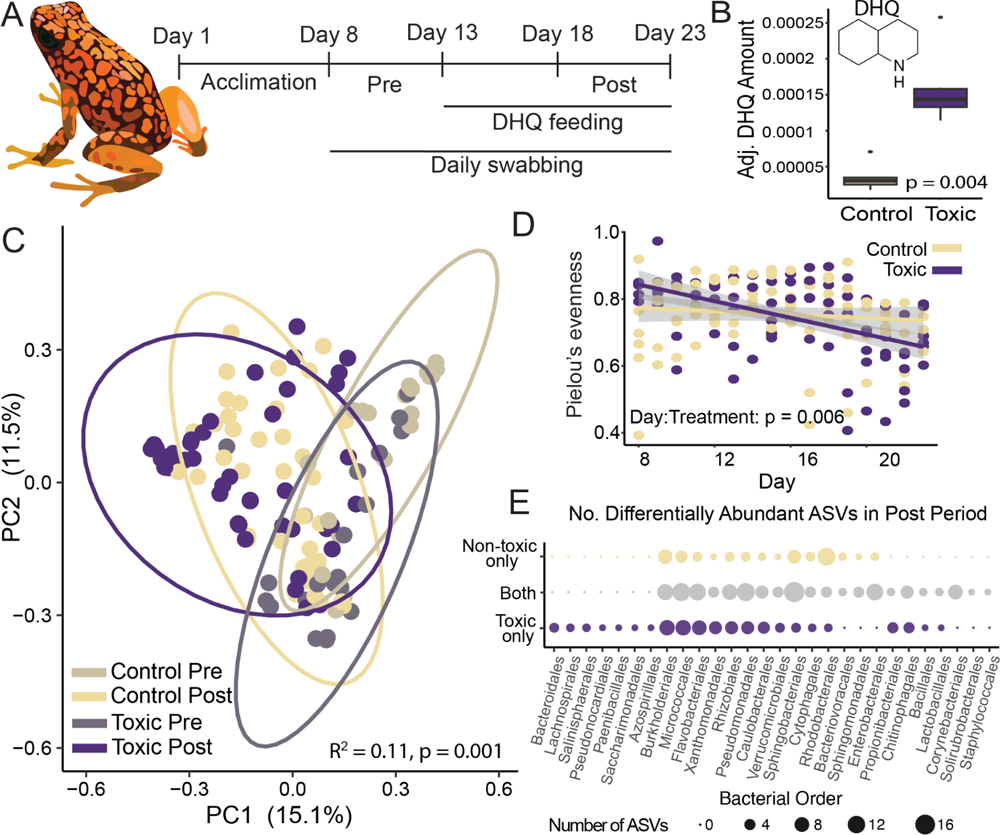
Bacterial communities shift in response to a single alkaloid. **(A)** Feeding experiment overview. **(B)** Quantification of DHQ shows accumulation in toxic frogs but not in control frogs. **(C)** Microbial communities shift in response to DHQ exposure using Bray-Curtis Dissimilarity. **(D)** Microbial communities on toxic frogs become less even over time. **(E)** Number of differentially abundant ASVs, as identified with ANCOMBC2, in post-treatment periods for DHQ fed frogs, control frogs, or in both treatments, grouped by bacterial order.

Both the bacterial and fungal communities changed in response to toxin feeding, although the fungal communities were less impacted (PERMANOVA: bacteria: Figure 3C, R^2^ = 0.11, p = 0.001, fungi: Figure 4A, R^2^ = 0.06, p = 0.001). The beta dispersion of the fungal communities decreased significantly in the post-DHQ feeding period for both the toxic and non-toxic frogs, suggesting that the differences in the PERMANOVA were due to a contraction of variation in the communities when disturbed (Figure 4B). There were shifts in microbial communities both in frogs fed DHQ as well as frogs not fed DHQ, likely due to changes resulting from the disturbance of daily swabbing. We observed a similar increase in species richness over time in both the toxic and non-toxic treatments (bacteria: R^2^ = 0.28, F_3,163_ = 22.59, Day:Treatment p = 0.97, fungi: Figure 4C, R^2^ = -0.015, F_3,107_ = 0.45, Day:Treatment p = 0.92). While the species richness changes did not differ across toxic and non-toxic treatments, the evenness of the bacterial communities decreased over time in the toxic *O. sylvatica* but not the non-toxic (bacteria: Figure 3D, R^2^ = 0.14, F_3,163_ = 9.7, Day:Treatment p = 0.006). This was not the case for the fungi, where there were no differences in community evenness across time or between treatments (Figure 4D, R^2^ = 0.02, F_3,107_ = 1.68, Day:Treatment p = 0.24).

**Figure 4.**
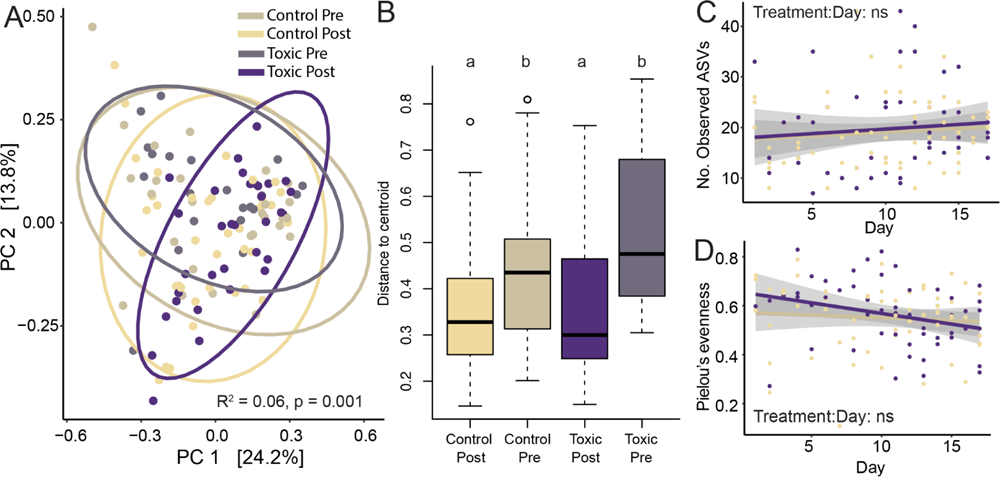
Fungal communities shift in response to a single alkaloid. **(A)** Fungal communities shift in response to DHQ exposure using Bray-Curtis Dissimilarity. **(B)** Shifts in fungal communities may be attributable to decreased distance to centroid in post-DHQ feeding communities. Letters correspond to statistical differences between groups based on a pairwise comparison of beta dispersion. **(C)** There are no significant differences between how the number of ASVs or **(D)** evenness change over time between toxic and non-toxic frogs.

Amplicon sequence variants (ASVs) that were differentially abundant between groups (toxic vs. non-toxic and pre vs. post) were identified with ANCOMBC 2 (Supplementary Table 2). Most differentially abundant taxa were identified in differences in structural zeros between groups. There were 134 ASVs that increased over time in both the toxic and non-toxic treatments, while 80 ASVs increased in only the toxic frogs and 54 increased in the non-toxic frogs (Figure 3E). Only one ASV decreased over time in the toxic group, while no ASVs decreased in the non-toxic group. In the fungi, there were fewer differentially abundant taxa. Four fungal ASVs were present in the post-period of toxic frogs and absent in the other groups. The fungal communities on the frogs were sparse and very different across frogs, therefore these taxa did not make up a significant portion of the total communities. Together, these results demonstrate that even a single alkaloid can alter the skin microbial community of a poison frog.

### Microbial isolates from a poison frog respond differentially to an alkaloid

Given that wild frog data combined with lab feeding experiment data suggested some bacterial and fungal taxa flourished in these toxic environments, we next tested how individual bacterial isolates responded to the presence of an individual alkaloid, DHQ. We isolated 132 strains from lab-reared non-toxic *O. sylvatica* across four media types. Based on full length 16S sequencing, these strains spanned 27 genera and 44 species of bacteria. Of these 27 genera, 22 were found in amplicon sequencing data from both the lab frogs and the frogs from the wild, suggesting that this strain library was representative of strains that occur on the frogs in the wild (Fig 5A). Using one strain per species, as identified by the closest full 16S BLAST match, we quantified the growth of these isolated strains in various DHQ concentrations (0%, 0.01, 0.1, 0.2, 1% w/v). Strains were categorized into three phenotypes based on growth (as measured by area under the growth curve) relative to growth without DHQ: enhanced, tolerant, and susceptible. Forty six percent of all tested strains were susceptible to DHQ (Fig 5B), 39% were tolerant to DHQ (Fig 5C) and 16% of strains were enhanced (Fig 5D, Supplementary Figure 3). Given the limitations of our data, it is difficult to directly compare strains to ASVs, although we saw concordance between growth with DHQ and differential abundance in the feeding experiment data at the genus level. For example, in the genera *Brachybacterium*, *Brevibacterium*, *Domibacillus*, and *Stenotrophomonas*, we cultured a strain that was enhanced with DHQ, and we observed ASVs in the lab amplicon sequencing dataset that were only present in frogs after DHQ feeding. While the strains were binned into general phenotype response categories, they exhibited a wide array of behaviors in growth with varying DHQ amounts, suggesting that across these strains, there may be a diverse set of strategies for dealing with DHQ, including potentially metabolizing DHQ for carbon to fuel additional growth. Overall, we were able to isolate multiple strains that are representative of the wild microbial community of poison frogs and determined that many of these strains grew unimpeded or even better with DHQ.

**Figure 5.**
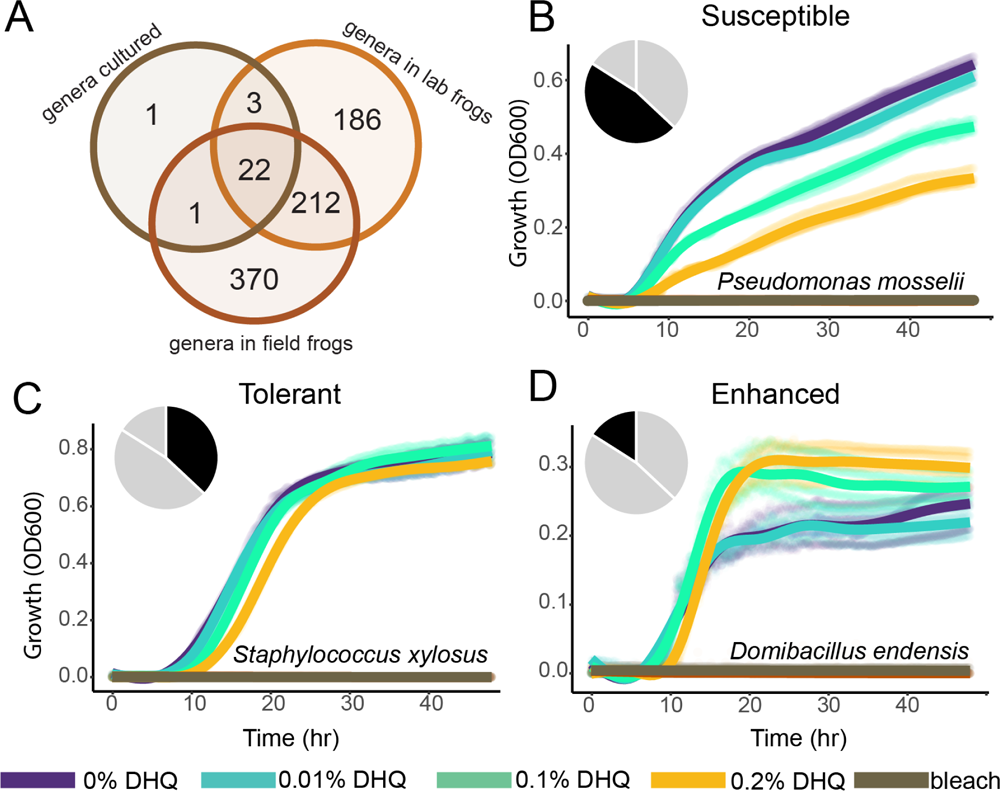
The majority of cultured bacterial strains are not affected by or grow better with an alkaloid. (A) Of the 27 bacterial genera cultured from the skin of *Oophaga sylvatica,* 22 are also found in amplicon sequencing data from both the lab experiment and the field. Growth patterns of strains can be categorized into those that are susceptible to DHQ (**B**; 45%), those that are tolerant (**C**; 39%), and those with enhanced growth in the presence of DHQ (**D**; 16%). Growth curves for all tested strains in Supplementary Figure 3.

### Some poison frog bacterial strains can incorporate carbon from decahydroquinoline

Given that several strains either grew better with the addition of DHQ or were able to be cultured on media that contained DHQ as the sole carbon source, we hypothesized that some of these strains were able to use DHQ as a carbon source. To test whether DHQ breakdown was occurring, we selected two bacterial strains that were isolated on media with DHQ as the sole carbon source, one from the genus *Providencia* (closest BLAST match *Providencia rettgeri*), and one from the genus *Serratia* (closest BLAST match *Serratia nematodiphila*), for isotope tracing experiments. The strains were first grown in ^13^C glucose to label the bacterial cells as much as possible, and then transferred to media with natural abundance DHQ dissolved in methanol (Fig 6A). Only methanol was added to the no-DHQ controls. Using nanoSIMS, the ^13^C atom percent enrichment (APE) was measured for samples taken at the beginning and end of the experiment (Fig 6B). We observed a decrease in ^13^C APE for all samples and treatments relative to time 0, suggesting that new carbon was incorporated by the bacteria in both treatments. However, *Providencia* sp. incorporated more new carbon in the DHQ treatment compared to the no-DHQ treatment (Fig 6C lower panel, R^2^ = 0.36, F_3,2860_ = 535.2, Treatment * Time p < 0.001). The results for *Serratia* are somewhat ambivalent because while there was a greater decrease in ^13^C enrichment in the DHQ treatment as compared to the methanol treatment (Fig 6C upper panel, R^2^ = 0.18, F_3,1074_ = 81.4, Treatment * Time p = 0.007), the ^15^N enrichments from ammonium were similar (Supplementary Fig 4A, R^2^ = 0.32, F_3,1074_ = 169.2, Treatment * Time p = 0.93), suggesting no difference in growth. Furthermore, the ^13^C enrichment for the DHQ treatment at time 0 was higher than for the no-DHQ treatment, which could inflate the differences observed across groups, but we do not have an explanation for why the time 0 isotope enrichments were different for the two treatments. We also observed that *Serratia* sp. incorporated significant quantities of ^15^N-ammonium in these experiments, while *Providencia* sp. uptake of ^15^N-ammonium was below detection in both treatments. Based on the ^13^C data (Fig 6C, Supplemental Fig 4B), *Providencia* sp. was growing, and if it was maintaining stoichiometry, it may have used the nitrogen from DHQ for growth instead of from the labeled ammonium. These results suggest that *Providencia* sp. was able to utilize DHQ as a carbon source for growth, and potentially as a nitrogen source.

**Figure 6.**
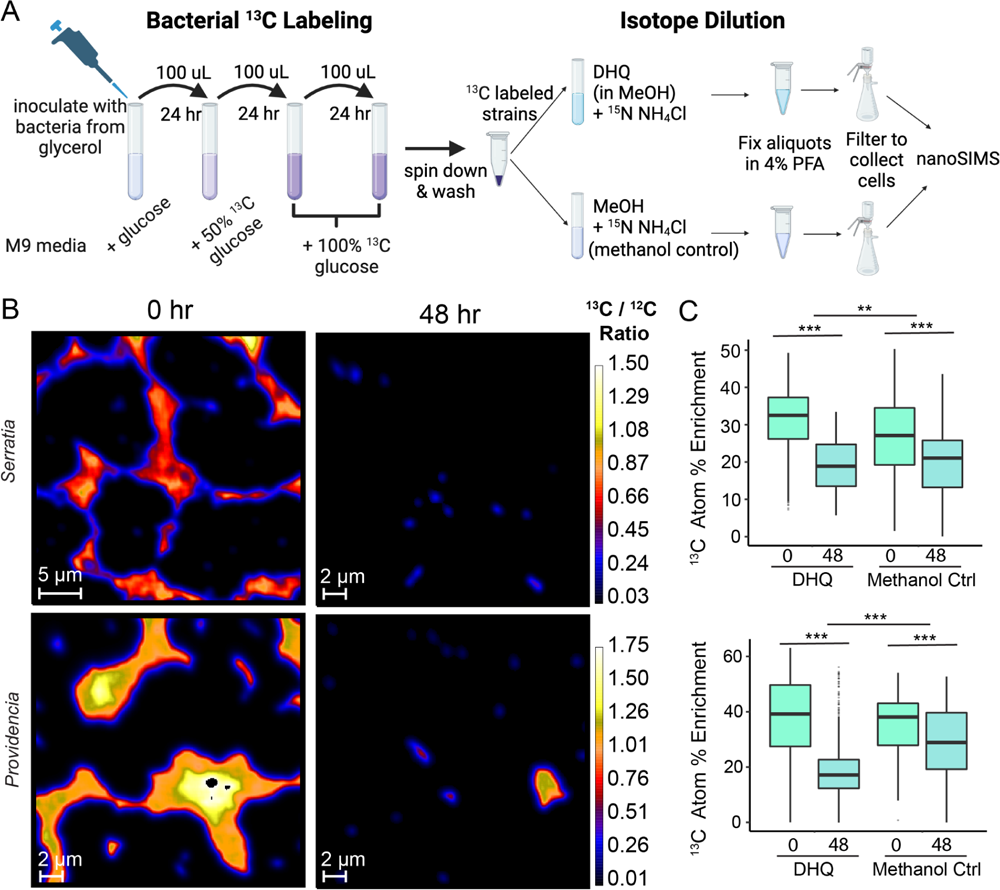
Reverse isotope tracing suggests bacterial metabolism of an alkaloid. **(A)** Poison frog strains were grown in 13C-glucose to label the cells with ^13^C and then they were assayed for incorporation of natural abundance DHQ carbon based on subsequent dilution of ^13^C. **(B)** For both *Serratia* sp. and *Providencia* sp., ^13^C/^12^C ratios as measured with nanoSIMS decrease over a 48-hour period. **(C)** ^13^C atom percent enrichment decreases significantly more across time in the DHQ treatment than in the control treatment with no DHQ. *** denotes p value < 0.001, ** denotes p value < 0.01.

## Discussion

This study explored how alkaloids shape the skin microbial communities of South American poison frogs, organisms with a long evolutionary history of alkaloid exposure. Contrary to our expectations, we found that poison frog species with high alkaloid loads had microbial communities that were more diverse and speciose than microbial communities of closely related, non-toxic frogs. Specifically, we found that the quantity of alkaloids, not the type or diversity of alkaloids, contributed to changes in community structure. One possible explanation for our results is that different alkaloid classes have promiscuous binding across microbial cell surface proteins, making the total alkaloid load stronger than alkaloid diversity when it comes to shaping the microbial community. This is supported by previous work showing that alkaloids across various classes of *Oophaga pumilio* poison frogs show conserved binding patterns in molecular binding assays against acetylcholine receptors ^32^, which supports the idea that there is promiscuity at the molecular level across different alkaloid types. It is possible that similar patterns hold for bacterial target proteins, where alkaloids across structural classes have promiscuous binding to microbial cell surface proteins. In this case, an increase in alkaloid diversity would not impact microbes as much as an increase in alkaloid load, which is what we observed in our study. Further work would be needed to identify microbial protein targets of poison frog alkaloids.

Our data corroborates the idea that location is an important driving factor in poison frog skin microbiome communities, and adds the critical axis of toxicity level to our understanding of community structure. In our study, and in many other studies of amphibian microbiomes, location has been identified as a major factor in determining microbial community composition ^33–35^. Another microbiome study on poison frogs found that between three species of dendrobatids collected in the same location, one chemically defended and two not, there were no differences in beta diversity, number of observed taxa, or phylogenetic diversity across the species ^31^. In this study the primary role of location in determining skin microbial community composition may have overpowered any differences in microbiomes due to alkaloids, which would explain why we observed a relationship between alkaloid load and microbial diversity where they did not. We sampled frogs from multiple locations, with each location including both toxic and non-toxic frogs, and found that microbial communities were indeed shaped by alkaloid load. The changes in microbial communities that we observed in this study are likely underestimates of the actual importance of these alkaloids in shaping the poison frog microbiome. From this data, it is difficult to disentangle the effects of the host species and geographically available microbes from the alkaloid-based differences in microbial communities. The relationship we observed in these results between higher alkaloid load, higher species richness, and lower evenness may occur because some microbial taxa that are highly abundant in non-toxic frogs may not grow well in the presence of alkaloids, which would provide opportunities for other, more rare taxa to proliferate. By comparing across many different locations and species, our work provides a more comprehensive and complex view of poison frog skin microbes, suggesting that there is an interaction between toxicity and microbial communities.

Beyond ecological correlations, we carried out the first experiment to directly measure how the skin microbiome of a poison frog changes when exposed to an alkaloid *in vivo*. These results corroborate our findings from the field, as we also identified a shift in microbial community and a diversification of rare taxa in frogs fed DHQ. We also showed variable growth responses of individual strains of bacteria isolated from *Oophaga sylvatica* to DHQ. While some of the isolated strains grew worse with DHQ, the majority exhibited no detectable effect or even increased growth in the presence of the alkaloid. Prior research examining the impact of poison frog alkaloids on microbes found alkaloids to be broadly antimicrobial ^28–30^, and decahydroquinoline alkaloids specifically have a documented antimicrobial activity against both bacteria and fungi ^28^. We have expanded on this work by looking beyond common lab microbial strains, focusing on the commensal microbiome of poison frogs. While Macfoy et al. and Mina et al. found alkaloids to be broadly antimicrobial against individual strains, we found that not only are there many strains from poison frogs that are not impacted by, or even grow better with an alkaloid, but also, the entire microbiome is made more diverse in the presence of alkaloids. Typically, acute exposure to compounds like antibiotics or pollutants make microbial communities less diverse ^36,37^. However, poison frogs have likely co-evolved with their microbial communities such that alkaloid-favoring taxa remain consistent members of the community, even when the frogs are not actively consuming alkaloids.

Finally, our nanoSIMS reverse isotope tracing experiments suggest that at least one strain was able to utilize DHQ as a partial carbon source for growth. The DHQ used in this experiment is the base ring structure of decahydroquinolines and is not a molecule found naturally on a frog. Regardless, the fact that these strains may be able to metabolize even small amounts of this DHQ supports the larger idea that there are strains on the skins of poison frogs that are adapted to not only survive in the presence of alkaloids, but also utilize alkaloids as a carbon source. In their cross-species comparison, Varela et al. found that the microbial communities from the toxic frog, *Dendrobates tinctorius*, were enriched for microbes with aromatic compound degradation capabilities ^31^. This function may be related to microbial metabolism of alkaloids. The microbial metabolism of alkaloids is not widely documented, which may be due to a lack of exploration of this phenotype or due to this phenotype truly being rare amongst microbes ^38^. Either way, this study provides multiple microbial candidates that seem to be able to metabolize these complex and ubiquitous molecules. Future studies could combine transcriptomics, genetics, and microbial engineering to understand these pathways and potentially utilize them for the metabolism of other alkaloids.

While alkaloid metabolism is relatively rare, it is unsurprising that it has arisen in this alkaloid rich environment. Similar microbial adaptations to toxic environments exist in other organisms, including herbivorous insects ^39–42^. This phenomenon also occurs in the gut microbiota of the desert woodrat, which is able to metabolize the tannins and oxalates from the creosote plant, reducing the toxic effect on the host ^43–45^. Similar to our results, exposure to a toxin that the woodrat host has experienced previously results in an increase in microbial community diversity, suggesting that this pattern may be generalizable across microbial communities in hosts that have evolved to consume high quantities of toxins. Animals that consume high quantities of toxic compounds may be reservoirs for bacteria and fungi with unique and useful metabolic pathways.

## Summary

This study is the first to characterize the impact of alkaloids on a poison frog host commensal microbiome of bacteria and fungi, and demonstrates that alkaloids play a significant role in structuring the microbial communities of poison frogs, potentially mediated by the alkaloid metabolic capabilities of strains. This work broadens our understanding of how long term exposure of a microbial community to exogenous compounds, in this case, alkaloids, alters microbial community composition and provides additional avenues for exploration of microbial metabolism of alkaloids and the impact of alkaloid-induced microbiome shifts on host health.

## Data availability

Raw data files, processed data, and data analysis files are available either as zip files accompanying the manuscript or on public repositories (pending acceptance). All mass spectrometry data (including GC/MS and LC/MS) can be found on DataDryad (pending acceptance). All 16S and ITS amplicon sequence FASTQ files can be found on SRA (pending acceptance). Full length 16S sequences from isolates are accessible on GenBank (pending acceptance). Bacterial and fungal strains are available from the authors upon request.

## Acknowledgements

We thank Courtney Swink and Christina Ramon for their assistance in conducting experiments at Lawrence Livermore National Laboratory. We thank the Relman lab for their input and suggestions on data analyses. We thank Tad Fukami for generously offering his lab space to perform the library preps and for guidance and suggestions about data analysis throughout. In Ecuador, Andrea Terán Valdéz helped with permit procedures and coordinating field work. We thank Jeff Bishop at the University of Oregon Genomics and Cell Characterization Facility for completing the library preps and sequencing for the feeding experiment samples. The authors acknowledge that this research was conducted on the ancestral lands of the Muwekma Ohlone people at Stanford. We understand the implications of the historical and present colonialism the Ohlone people experience and celebrate their continued stewardship of their lands.

## Funding

This work was supported by the New York Stem Cell Foundation (LAO). LAO is a New York Stem Cell Foundation – Robertson Investigator. SNC is supported by an NSF Graduate Research Fellowship (DGE-1656518) and a Stanford Graduate Fellowship in Science and Engineering. AAB is supported by a NSF Graduate Research Fellowship (DGE-1656518) and an HHMI Gilliam Fellowship (www.hhmi.org, GT13330). Work at Lawrence Livermore National Laboratory was conducted under contract DE-AC52-07NA27344 and funded by the microbiospheres Science Focus Area grant # SCW1039 by the US Department of Energy’s Office of Biological and Environmental Research. Part of this work was supported by the Vincent Coates Foundation Mass Spectrometry Laboratory, Stanford University Mass Spectrometry (RRID:SCR_017801), utilizing the Thermo Exploris 240 BioPharma LC/MS system (SCR_022216), and in part by NIH P30 CA124435, utilizing the Stanford Cancer Institute Proteomics/Mass Spectrometry Shared Resource.

## Author Contributions

Conceptualization: SNC, MMM, XM, LAO

Data Curation: SNC

Formal Analysis: SNC, MMM, TM, AAB

Funding Acquisition: LAO, XM, PKW, LAC

Investigation: SNC, MMM, TM, AAB, NAM, CV, EET

Methodology: SNC, MMM, XM, PKW, TM

Resources: XM, LAO

Supervision: MMM, LAO, LAC

Visualization: SNC

Writing - Draft Preparation: SNC

Writing: Review & Editing: AAB, XM, PKW, CV, NAM, TM, LAO, LAC, MMM

## Declaration of Interests

The authors declare no competing interests.

## Methods

### Field collection of frogs

Individuals from 11 species of dendrobatid frogs (*Ameerega bilinguis*, *Ameerega parvula*, *Oophaga sylvatica*, *Epipedobates tricolor*, *Epipedobates boulengeri*, *Epipedobates darwinwallacei*, *Hyloxalus toachi*, *Hyloxalus infraguttatus*, *Allobates zaparo*, *Allobates talamancae*, and *Allobates femoralis*) were collected from 9 locations throughout Ecuador (Fig 1A). Animals were caught during the day and stored individually in moist plastic bags until the evening when frogs were rinsed with sterile water and swabbed 10 times (up and back is once) with sterile cotton swabs (Puritan 25-806 2WC). After swabbing, frogs were anesthetized with topical application of benzocaine and euthanized by cervical transection. Swabs were immediately placed into 100% ethanol and kept at −20 °C for long-term storage. Half of the dorsal skin from each frog was stored in methanol for alkaloid analyses until shipment to the US, wherein they were stored at −20 °C. Collections and exportation of specimens were done under permits (No. 0013-18 IC-FAU-DNB/MA; Export permit: No. 214-2019-EXP-CM-FAU-DNB/MA; CITES export permit No. 19EC000036/VS) issued by the Ministerio de Ambiente de Ecuador. All procedures were approved by the Administrative Panel on Laboratory Animal Care Committee of Stanford University (Protocol 34153).

### Alkaloid extraction and quantification

Skins were processed for alkaloid extractions as described in Alvarez-Buylla et al, 2023 ^46^. Briefly, the methanol in which the skin was stored was filtered and spiked with (-)-nicotine (Sigma Aldrich, N3876-100ML), then stored at −80 °C for 24 hours to precipitate lipids and proteins. After filtering into new vials, a 100 µL aliquot was added to a gas chromatography / mass spectrometry (GC/MS) autosampler vial. GC/MS analysis was performed on a Shimadzu GCMS-QP2020 instrument with a Shimadzu 30 m x 0.25 mmID SH-Rxi-5Sil MS column closely following the protocol outlined in Saporito et al, 2010 ^47^. In brief, separation of alkaloids was achieved with helium as the carrier gas (flow rate: 1 mL/min) using a temperature program from 100 to 280 °C at a rate of 10 °C/min. This was followed by a 2 min hold and additional ramp to 320 °C at a rate of 10 °C/min for column protection reasons, and no alkaloids appeared during this part of the method. Compounds were analyzed with electron impact-mass spectrometry (EI-MS). The GC-MS data files were exported as CDF files and the Global Natural Products Social Molecular Networking (GNPS) software was used to perform the deconvolution and library searching against the AMDIS (NIST) database to identify all compounds ^48^. For deconvolution (identification of peaks and abundance estimates) the default parameters were used. Through the deconvolution process, molecular features were reported as rows/observations, while m/z intensities were reported as columns/variables. Automatic library search was obtained from reference libraries of natural products (NIST, Wiley, University of CORSICA, GNPS), and our resulting dataset was filtered to keep only our nicotine standard and alkaloids previously found in poison frogs from the Daly 2005 database, or compounds with the same base ring structure and R group positions as the classes defined in Daly 2005 ^49^. Once the feature table from the GNPS deconvolution was filtered to include only poison frog alkaloids and nicotine, the abundance values (ion counts) were normalized by dividing by the nicotine standard and skin weight, and we included the 10 top database hits in our library search. The resulting filtered and normalized feature table was used for all further analyses and visualizations. All normalization steps were carried out with R (version 4.0.4).

### DNA extraction from field swabs

Swabs in ethanol were stored at −20°C until DNA extraction. DNA was extracted from swabs using the Qiagen PowerSoil Pro Kit, adapted for use with swabs stored in ethanol. Briefly, swabs were quickly (2-3 s) vortexed in ethanol and dried at 37 °C until dry (approximately 2 hours). The remaining ethanol was centrifuged at 15294 rcf for 15 min. The ethanol was discarded, and the pellet was washed once with 200 µL PBS, then centrifuged at 15294 rcf for 5 min. After removal of PBS, 800 µL of lysis buffer (CD1) was added to the pellet and allowed to incubate at room temperature for 10 min, after which the solution was transferred, along with the dried swab, to the PowerBead Pro Tube. Swabs were included through the addition of solution CD2, at which point swabs were removed and the protocol continued per manufacturer instructions. Solution C6 was allowed to incubate for 10 min prior to centrifugation. Extracted DNA was stored at 20 °C for prolonged storage, and 4 °C when in use.

### Library preparation and sequencing of field swabs

For amplification of the taxonomy-inferring 16S gene for bacteria and ITS gene for fungi, the V4 and ITS1 regions were amplified and sequenced. Samples were randomly distributed across 96 well plates. In each well, 4.2 µL of extracted DNA was combined with 5 µL of MyTaq Red polymerase (Bioline, Meridian Bioscience), 0.4 µL of 10 uM forward primer (nexF-N_3-6_-515f for 16S or nexF-N_3-6_-ITS1f-KYO1 for ITS), and 0.4 µL of 10 uM reverse primer (nexF-N_3-6_-806r for 16S or nexF-N_3-6_-ITS2-KYO2 for ITS) (35,36). 16S PCR parameters were as follows: denaturation at 95 °C for 2 min followed by 35 cycles of denaturation at 95 °C for 20 s, annealing at 52.5 °C for 20 s, and extension at 72 °C for 50 s with a final extension of 10 min at 72 °C. The same protocol was used for ITS amplification with 50 °C used as the annealing temperature. Amplification of PCR 1 product and lack of amplification of no DNA controls was confirmed from 12 randomly selected samples per 96 well plate using the Lonza FlashGel^TM^ system (Lonza, 57067) with the FlashGel^TM^ DNA Marker Ladder (Lonza, 50473) for size confirmation. Samples were subsequently barcoded in the second PCR using unique combinations of P5-Hamady-nexF and P7-Hamady-nexR primers ^50^. For this PCR, 1 µL of template DNA from PCR 1 was added to 5 µL of MyTaq Red, 3.2 µL of water, and 0.8 µL of 10 uM primer (combination of forward and reverse). Samples were amplified with the following protocol: denaturation at 95 °C for 2 min followed by 8 cycles of denaturation at 95 °C for 20 s, annealing at 50 °C for 20 s, and extension at 72 °C for 50 s, followed by a final extension of 10 min at 72 °C. PCR products were cleaned and size selected with AMPure Beads in a ratio of sample to beads of 1:1.2. All 16S libraries were pooled in equal volumes and all ITS libraries were separately pooled in equal volumes. Pooled libraries were run on an Agilent TapeStation to confirm adequate size distribution and quantified with Qubit. Samples were sequenced by Azenta Life Sciences (Burlington, MA, USA) with a 2x250 Illumina MiSeq configuration.

### Feeding experiment

Lab-raised Diablito frogs (*Oophaga sylvatica,* N=12) were purchased from Indoor Ecosystems (Whitehouse, OH, USA). During the experiment, frogs were housed individually in plastic containers (Sterilite 16428012) with moist, autoclaved paper towels and plastic reptile hides (XYZ Reptiles, Palmetto Bay, FL, USA). Paper towels were replaced daily and sprayed with new sterile water to maintain humidity in the boxes. Frogs were fed ∼ ⅛ teaspoon of fruit flies dusted with calcium powder (Repashy Calcium Plus) and vitamin powder (RepCal Herptivite) daily during the experiment. Frogs were split into two groups: those fed the alkaloid decahydroquinoline (DHQ, Santa Cruz Biotechnology, Santa Cruz CA, USA, N = 6) and those fed a water vehicle control (N = 6). Frogs from the same group housing tanks were split between two treatment groups, and sex was roughly evenly distributed across the two groups. Frogs were allowed to acclimate to their new housing for one week, after which (Day 8 and on) they were swabbed daily for the remainder of the experiment. For swabbing, frogs were rinsed with sterile water and then swabbed 10 times (up and back is once) on the dorsal side using Puritan Sterile Tipped Polyester Applicators (Puritan Medical Products, 25-206 1PD BT). Alkaloid feeding began 5 days after the onset of swabbing (Day 13) and continued for the duration of the experiment, for a total of 10 days. Frogs fed decahydroquinoline (DHQ) were fed 10 µL of a 0.01% DHQ in water solution daily via pipetting directly into the mouth. Control frogs were fed 10 µL of water in the same manner daily. Swabbing occurred prior to alkaloid feeding on days when both occurred. Swabs were immediately placed on ice after collection and subsequently stored at −20 °C until DNA extraction. All procedures were approved by Administrative Panel on Laboratory Animal Care Committee of Stanford University (protocol #33691).

### DNA extraction, library prep and sequencing for feeding experiment samples

DNA was extracted from swabs using the Qiagen PowerSoil Pro Kit, adapted for use with dry swabs. Swabs were included through the addition of solution CD2, at which point swabs were removed and the protocol continued per manufacturer instructions. Extracted DNA was sent to the University of Oregon Genomics and Cell Characterization Core Facility for library preparation and sequencing. The 16S V4 region was amplified using 515F (GTGYCAGCMGCCGCGGTAA) and 806R (GGACTACNVGGGTWTCTAAT) primers (35). The ITS region was amplified using the ITS1F-kabir (CTTGGTCATTTAGAGGAAGTAA) and the ITS2R-kabir (GCTGCGTTCTTCATCGATGC) primers (38,39). Samples were sequenced in a 2x300 configuration on Illumina MiSeq v3.

### Analysis of 16S and ITS amplicon sequencing

Both the field sample and feeding experiment 16S and ITS datasets were cleaned and processed with the R package dada2 (version 1.26.0) ^51^. Taxonomy was assigned to 16S reads with the Silva NR99 version 138.1 dataset, and to ITS reads with the UNITE database (versions 10.05.2021 for the field data and 29.11.2022 for the feeding experiment data). The amplicon sequence variants (ASVs) generated from dada2 were incorporated into phyloseq objects for additional data analysis (phyloseq version 1.42.0) ^52^. As a conservative measure, any ASVs found in no DNA control sequencing reactions were removed from the datasets. For the 16S datasets, any ASVs with a phylum designation that was uncharacterized, “NA”, or Eukaryotic, or a family designation of Mitochondria were removed from the datasets. For the ITS data, uncharacterized and “NA” phyla were removed. To account for differences in sampling depth, alpha diversity analyses were conducted with rarefied datasets. For 16S alpha diversity comparisons in the field samples, samples with fewer than 10,000 reads were excluded, and then all samples were rarefied to 90% of the read number of the sample with the fewest reads. Because the number of reads were lower from the feeding experiment samples, samples from the feeding experiment with fewer than 5,000 reads were excluded, and then all samples were rarefied down to 95% of the read number of the sample with the fewest reads to maintain as much data as possible. For beta diversity-based analyses in all datasets, any ASVs found in only one frog were removed from the full, non-rarefied dataset, and counts were converted into relative abundance. In the field data, ASVs were agglomerated at the genus level to facilitate comparison across the disparate ASVs found across the poison frog species. Bray-curtis dissimilarity was used for multidimensional scaling using Principal Coordinates Analysis (PCoA). Differential abundance analyses were conducted with ANCOMBC2 (version 2.0.3). Statistical analyses were performed in R (version 4.2.1) with vegan (version 2.6.4), pairwiseAdonis, dendextend (version 1.17.1), and pvclust (version 2.2.0).

### Feeding experiment alkaloid quantification

At the conclusion of the laboratory feeding experiment, frogs were anesthetized with benzocaine application to the ventral side and euthanized via cervical transections. Dorsal skins were removed and placed in Trizol (Invitrogen) and immediately homogenized in BeadBug tubes for 6 cycles of 30s at speed 5 with a 1 min delay between cycles, then stored at −80 °C until used for alkaloid extractions. Homogenized samples were thawed and weighed to account for size of skin used for analysis. The liquid (∼1 mL) was removed from the BeadBug tubes and transferred to new tubes with 150 µL of chloroform. Tubes were shaken for 15 s, then left at room temperature for 3 min. Samples were then spun down at 15682 rcf for 15 min at 4 °C. The top layer was removed and 300 µL of 100% ethanol was added to the remaining organic phase, then mixed by inverting 10 times and allowed to settle for 3 min at room temperature. The samples were centrifuged at 2000 rcf for 5 min at 4 °C to pellet the DNA, then 300 µL of the supernatant was transferred to a new tube along with 900 µL of acetone. The samples were mixed by inversion for 10-15 s, then incubated for 10 min at room temperature. The samples were then centrifuged for 10 min at top speed at 4 °C to pellet the protein precipitate, then 1 mL of the supernatant was removed and placed into glass vials. The supernatant was filtered through a 0.1 µM filter, then dried down under N_2_ gas and reconstituted in 270 µL methanol with 30 µL of a 700 µM nicotine solution to be used as an internal standard. The samples were centrifuged at 5000 rcf for 5 min to pellet any remaining particulate, and 100 µL was removed and placed into autosampler vials for LC/MS analysis. Samples were diluted 1:5 to fit within the dynamic range of the instrument. The samples were analyzed by ESI-MS on the Waters Acquity UPLC and Thermo Exploris 240 BioPharma orbitrap mass spectrometer. A Sequant zic-HILIC 3.5u 100 x 2.1 mm LC column was used with a flow rate of 02 mL/min with an injection volume of 5 µL. The solvents used were 5 mM ammonium acetate and 0.1% formic acid in water (Solvent A) and 0.1% formic acid in acetonitrile (Solvent B). The solvents were used with the following gradient: 10% A, 90% B for 0.5 min, ramped to 20% B in 14 min and held for 2 min. Full Scan MS and targeted Full Scan MS2 data were acquired: FullMS mass range 100-1000 with resolution 120,000; FullMS2 of m/z 140.1434 (DHQ) and FullMS2 of m/z 163.123 (nicotine ISTD) isolation window = 2, microscans = 1, with fixed absolute HCD collision energy of 20 and resolution 15,000. To calculate the adjusted DHQ amounts, the area under the curve (AUC) for DHQ was divided by the AUC for nicotine, then by the calculated skin weight for each sample.

### Microbial isolate library generation

Five microbial isolate libraries were made using five media types; each originated from a swab from a single laboratory reared, non-toxic *Oophaga sylvatica* individual (N=5 total) that had not been used in the feeding experiments. Frogs were rinsed with sterile water and swabbed 10 times (up and back is once) on the dorsal side. Swabs were immediately struck out onto solid media plates. Multiple media types were used to encourage higher diversity of cultivated strains. The five media types used included 1% tryptone media (tryptone: Fisher BioReagents, Fisher Scientific), R2A (BD Difco, Fisher Scientific), LB (BD Difco, Fisher Scientific), tryptic soy agar with sheep’s blood (pre-made plates: Thermo Scientific; tryptic soy agar: BD Difco, Fisher Scientific; defibrinated sheep’s blood: Thermo Scientific, Fisher Scientific), and M9 minimal media (BD Difco, Fisher Scientific) with 0.01% w/v decahydroquinoline (Santa Cruz Biotechnology) as the sole carbon source. Plates were sealed with Parafilm and incubated at 21°C. When colonies were large enough to be picked (ranging from about 12-168 hours), they were isolated three times by re-streaking onto new plates of the same media type. A single colony from the third plate was used for both strain identification and glycerol stock production. To identify colonies, samples were prepared for full length 16S sequencing or ITS region 1 Sanger sequencing as follows. Individual colonies were placed into 10 µL of sterile water and 1 µL of this input was used in a PCR reaction with 1 µL each of forward primer 27F (5’-AGAGTTTGATCMTGGCTCAG-3’) and reverse primer 1492R (5’-TACGGYTACCTTGTTAYGACTT-3’) ^53^ for 16S or forward primer ITS4 (5’-TCCTCCGCTTATTGATATGC-3’) and reverse primer gITS7 (5’-GTGARTATCGARTCTTTG-3) for ITS ^54,55^, 12.5 µL of OneTaq polymerase (New England Biolabs) and 9 µL water. Bacterial PCR parameters were as follows: denaturation for 30 s at 94°C followed by 30 cycles of denaturation for 15 s at 94 °C, annealing for 30 s at 55 °C and extension for 2 min at 65 °C, then a final extension period of 5 min at 65 °C. Fungal PCR parameters were as follows: denaturation for 30 s at 94 °C followed by 30 cycles of denaturation for 15 s at 94 °C, annealing for 30 s at 55 °C and extension for 2 min at 68 °C, then a final extension period of 5 min at 68 °C. Colony amplification was confirmed with gel electrophoresis, after which samples were cleaned with the E.Z.N.A Cycle Pure Kit (Omega). If no amplification occurred, DNA was extracted from colonies using the Qiagen Blood and Tissue Kit adapted for samples from gram positive bacteria. Cleaned DNA samples were submitted to Azenta for Sanger sequencing. Forward and reverse reads were aligned using the Geneious Alignment in Geneious Prime with the following parameters: global alignment with free end gaps, 65% similarity (5.0/-4.0) cost matrix, gap open penalty 12, gap extension penalty 3. The aligned sequences were BLASTed with Megablast on Geneious Prime to either the 16S ribosomal RNA database for bacteria or the nucleotide collection nr/nt for fungi. Glycerol stocks were made by combining equal parts of a sterile-filtered 50% glycerol and water solution with cultures grown overnight in liquid media matching the initial isolation media. Glycerol stocks were maintained in duplicate at −80 °C for downstream use.

### Quantification of microbial growth with an alkaloid

To measure the effect of the alkaloid DHQ on the growth of microbes isolated from *O. sylvatica*, individual strains were grown in five concentrations of DHQ: 0%, 0.01% w/v, 0.1% w/v, 0.2% w/v and 1% w/v in 1% tryptone media. DHQ stocks were maintained in methanol, so methanol volumes were kept consistent across treatments and were below 1% of the total solution volume. Strains were grown from glycerol stocks overnight at room temperature, shaking at 200 rpm in 1% tryptone then diluted to 10^6^ as determined from OD600 readings. Then, 180 uL of media and 20 uL of the diluted strain were added to each well. Plates were sealed with Breathe Easy (Sigma-Aldrich) membranes. Data was generated by measuring OD600 for four replicates of each condition at 5 min intervals for 48 hours on a BioTek Epoch2 plate reader for most strains, although slower growing strains were grown for 96 hours with reads every 8 min. Control wells were included for each DHQ concentration, and the average value for each control time point was subtracted from the experimental measurements for the respective treatments and time points. Growth curves were plotted using the plater package (version 1.0.4) and growthcurver (version 0.3.1) was used to fit curves and calculate the area under the curve. Areas under the curve were compared using ANOVA with a Tukey HSD post-hoc test. All statistical analyses were conducted in R (version 4.2.1). Strains with at least one treatment with statistically higher AUC than the control (no DHQ) were characterized as enhanced. Strains with at least one treatment with statistically lower AUC than control were characterized as susceptible, and strains with no significant differences in AUC were characterized as tolerant.

### Reverse labeling nanoSIMS experiment

Two bacterial strains, *Providencia* sp. and *Serratia* sp., that were cultured from *Oophaga sylvatica* skins using M9 media with DHQ as the sole carbon source were selected for reverse labeling isotope tracing analysis with nanoSIMS to test for metabolism of DHQ. The idea was to incubate ^13^C-labeled cells in unlabeled DHQ and examine the dilution of isotope over time, which would be consistent with utilization of DHQ carbon. Strains were grown from glycerol stocks in M9 minimal media with glucose overnight. In an attempt to fully label the cells with ^13^C, 100 µL of each strain was transferred from the unlabeled glucose M9 media to 5 mL of M9 media with 50% ^13^C glucose (99% ^13^C, Cambridge Isotope Laboratory) and 50% natural abundance glucose and allowed to grow overnight at room temperature, shaking at 210 rpm. Then 100 µL of each strain was transferred to 5 mL of 99% ^13^C glucose twice. After the second round of growth in 99% ^13^C glucose, the strains were spun down for 15 min at 3000 rcf to pellet the cells, then washed twice with PBS to remove residual glucose. After the second wash, the strains were resuspended in PBS and divided amongst one of three treatments: experimental, methanol control, or kill control. In the experimental condition, 100 µL of resuspended cells were added to M9 media with decahydroquinoline (DHQ; C_9_H_17_N; natural carbon abundance: 99.93% ^12^C, 1.07% ^13^C) dissolved in methanol and ^15^NH_4_Cl. In the methanol control, the strains were added to M9 media with an equivalent amount of methanol and ^15^NH_4_Cl as used in the experimental treatment, but with no DHQ. For the kill control, 100 µL of the resuspended cells were fixed in paraformaldehyde (PFA) at a final concentration of 4% for two hours, then washed once with PBS and added to M9 media with DHQ and ^15^NH_4_Cl, as in the experimental treatment (Figure 4A). In addition to these treatments, a no-addition control was generated to determine the 13C baseline for the strains prepared in natural abundance glucose and subsequently transferred to M9 media with DHQ dissolved in methanol with unlabeled NH_4_Cl. Initial time point (t0) samples were collected immediately after inoculation. At 48 hours (t48), the unfixed samples (experimental, methanol control and no addition control) were fixed with PFA at a final concentration of 4% for two hours, then washed once and resuspended in 1 mL of PBS. The kill control was spun down at 3000 rcf for 15 min to pellet the cells, then the supernatant was removed and the pellet was resuspended in PBS.

Samples were prepared for nanoSIMS following previously established protocols ^56^. Briefly, the t0 and t48 time points were filtered onto 0.2 µM pore white polycarbonate membrane filters (Whatman Nucleopore, GE Healthcare Life Sciences, Pittsburg, PA). Initial time points (t0) were undiluted while t48 samples were diluted 1:10 to adjust for higher cell concentration. Filters were rinsed with milliQ water, then left to dry completely. After drying, sterile scissors were used to cut slices of each filter and adhered to an analysis bullet using conductive tabs and sputter coated with ∼5 nm of gold. Isotope imaging was performed with a Cameca NanoSIMS 50 at Lawrence Livermore National Laboratory. A primary 133 CS + ion beam (2 pA, 150 nm diameter, 16 keV) was rastered over a 25 x 25 um analysis area with 256 x 256 pixels and a dwell time of 1 ms/pixel for 25-30 cycles. Prior to the analysis, areas were sputtered with 90 pA of Cs+ current to reach sputter equilibrium. Secondary ion images were simultaneously collected for ^12^C_2_^-^, ^12^C^13^C^-^, ^12^C^14^N^-^, ^12^C^15^N^-^ and ^32^S^-^ on individual electron multipliers in pulse counting mode, as described by Pett-Ridge and Weber ^57^.

All images were processed using L’image (version 11-1-2021, Larry Nittler, Carnegie Institution of Washington). Ion image data were corrected for deadtime and image shift across cycles before producing ^13^C^12^C/^12^C_2_ and ^12^C^15^N/^12^C^14^N ratio images. Regions of interest (ROIs) were autodetected in L’image using the ^12^C^14^N^-^ images. Ratios were averaged across cycles. Background corrections were performed for carbon ratios to adjust for differences across runs, and both carbon ratios and nitrogen ratios were converted to atom percent enrichment (APE) values based on the initial, pre-labeling (Ri) and final, post-labeling (Rf) ratios: APE = [Rf/(Rf + 1) – Ri/(Ri + 1)] · 100% ^58^. Difference-in-differences analyses were performed in R to compare the change in isotope enrichment between time 0 and 48 across the DHQ and methanol control treatments.

## Notes

### Competing Interest Statement

The authors have declared no competing interest.

