## Supplemental Figures 1-4 for "A toxic environment selects for specialist microbiome in poison frogs"

**Supplementary Figure 1. Number of individual toxins found on a frog does not correlate with square root transformed bacterial ASV counts.**

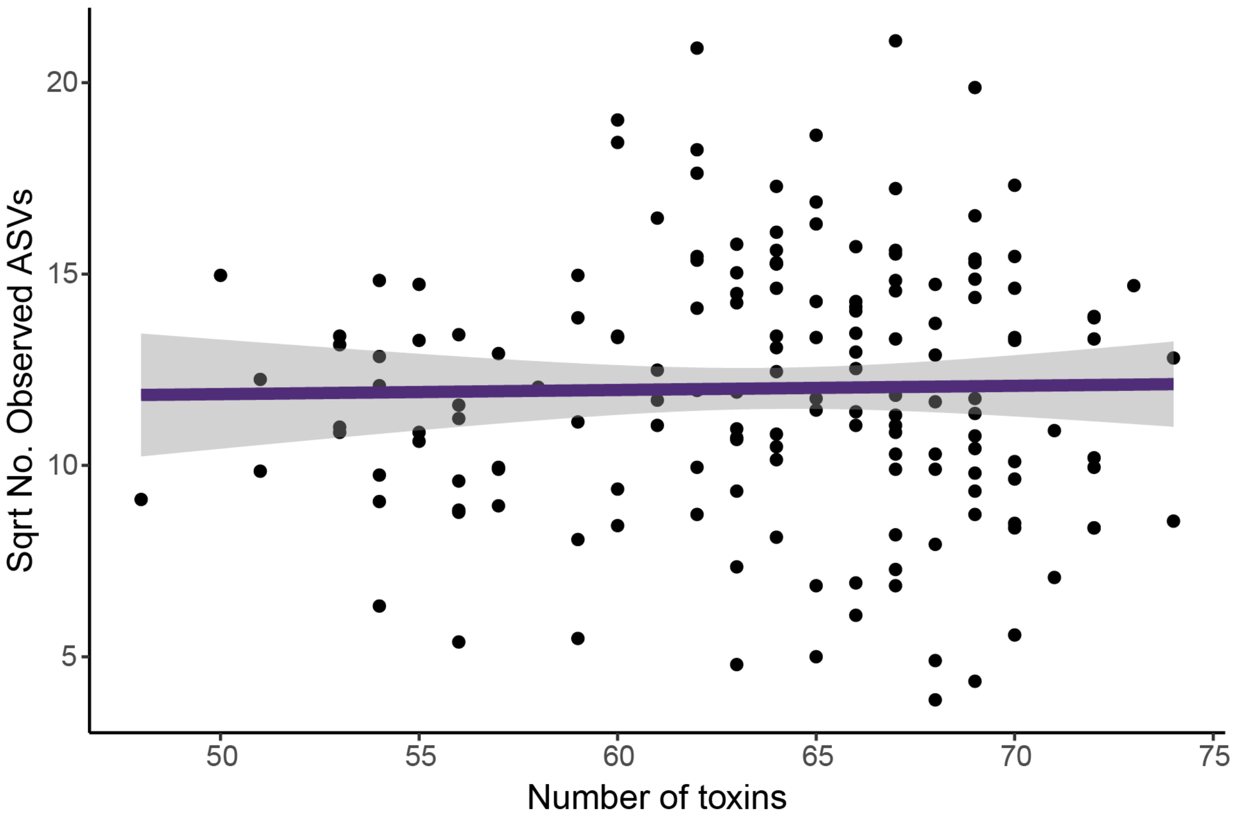


Each dot represents an individual frog. Number of toxins represents the number of individual alkaloids found in any quantity above zero. There is no significant linear relationship between number of toxins and square root transformed count of ASVs (R2 = -0.006, $F_{1,159}$ = 0.05, p = 0.82).

**Supplementary Figure 2. Decahydroquinolines are common alkaloids found on poison frogs, including *Oophaga sylvatica*.**

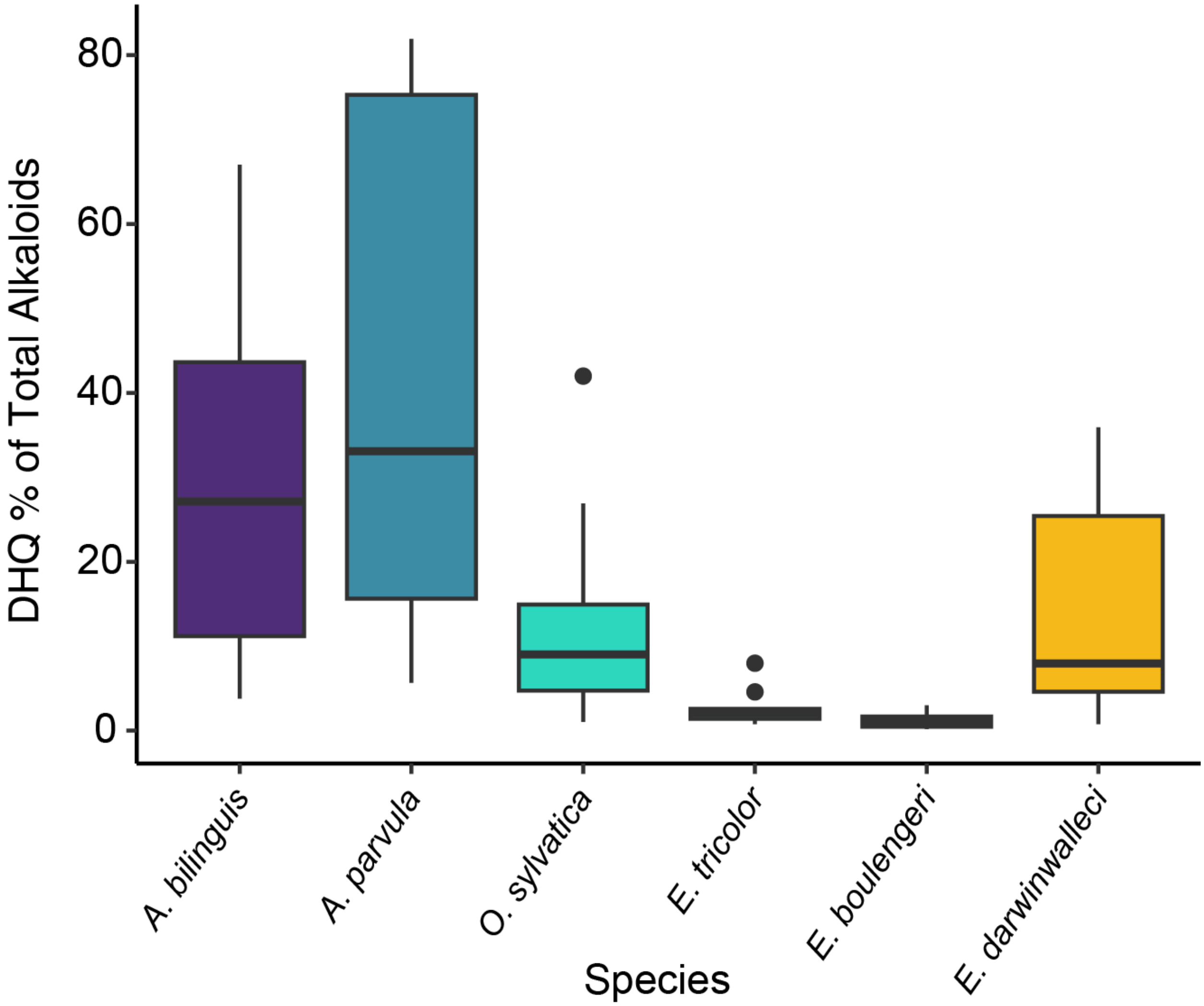


The 16 compounds that were categorized within the family decahydroquinoline from the NIST database were included. Dots are outlier points.

**Supplementary Figure 3. Growth curves from all characterized strains.**

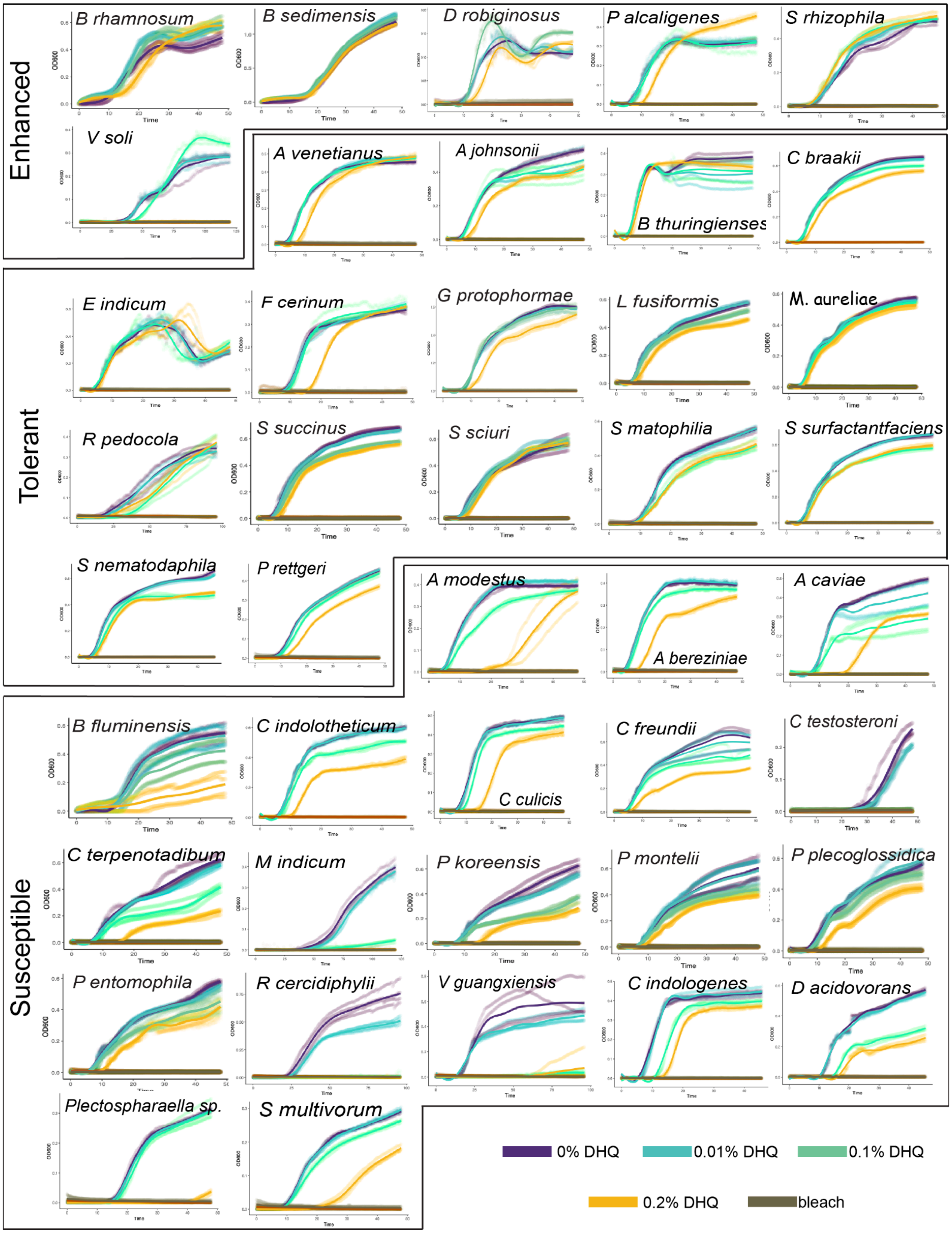

Growth was measured by OD600 values over time. N=4 replicates per condition. Strains were categorized as follows: enhanced when at least one DHQ condition had a statistically greater area under the curve than the 0% DHQ condition, tolerant when DHQ conditions were not different from 0% DHQ, and susceptible otherwise.

**Supplementary Figure 4**. **Nitrogen incorporation by strains and control enrichment patterns measured by nanoSIMS.**

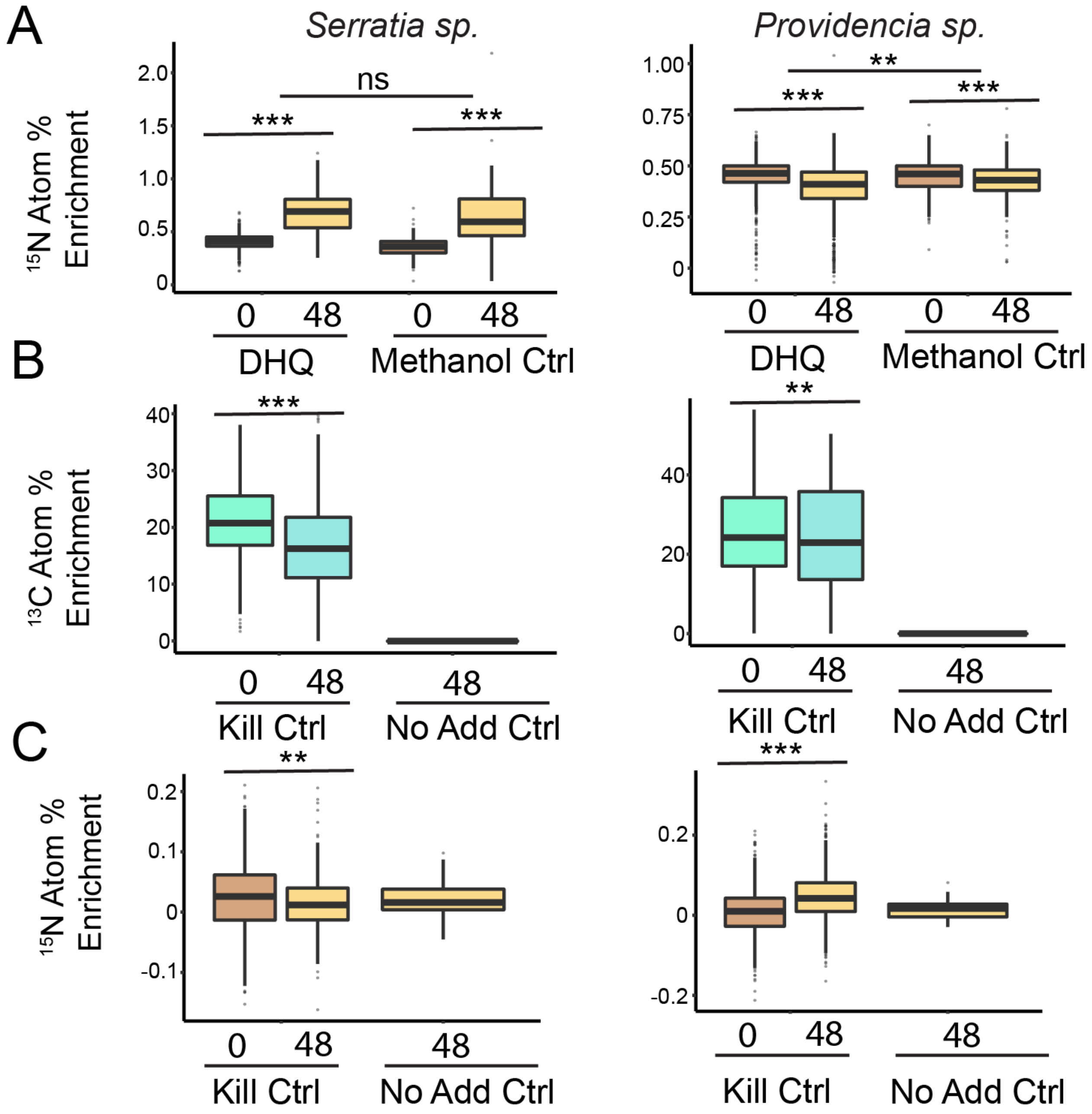


(A) *Serratia* strains incorporated $^{15}N$ from ammonium, demonstrating growth. *Providencia* did not incorporate $^{15}N$ from ammonium, suggesting that the *Providencia* that grew in the DHQ control preferentially utilized nitrogen from DHQ. (B) $^{13}C$ atom percent enrichment in control groups: kill controls were fixed with PFA prior to incubation with DHQ, no-add controls never received any labeled isotopes. (C) $^{15}N$ atom percent enrichment in control groups.
